# Harnessing Alpine Lake Bacteria for the Development of Synthetic Communities with Broad-Spectrum Antiviral Activity

**DOI:** 10.1101/2025.10.31.685752

**Authors:** L. Daniela Morales, Jessica Aubouy, Shannon David, Paul Seguin, Olivier Schumpp, Anna Carratalà

## Abstract

In aquatic ecosystems, phagotrophic protists and bacteriophages are major biological factors shaping bacterial populations. However, the extreme environmental conditions and low nutrient concentrations in alpine lakes limit the abundance of protists. Thus, bacteriophages may represent the main biological factor shaping bacterial communities in these ecosystems. We hypothesized that alpine lake bacteria populations harbor diverse antiviral strategies against bacteriophages that could be used to fight against other viral pathogens. To evaluate the presence of bacteria with broad-spectrum antiviral activity in high altitude Swiss lakes, we developed a sampling campaign in 24 alpine lakes in Switzerland. We assessed the decay of two human viruses (echovirus 11 and influenza A virus) in lakewater and characterized the microbial abundance and metabolic diversity in the collected samples. In addition, we obtained a collection of 223 bacterial isolates and selected bacteriophage resistant bacteria to build synthetic communities with enhanced antiviral activity against the two tested human viruses, and one plant virus of commercial significance (Potato Virus X). This confirmed our hypothesis and drew for the first time a relation between bacteriophage resistance and viral inactivation. Our findings emphasize the importance of alpine lakes as hotspots of microbial diversity with unexplored biotechnological applications.

## 1. INTRODUCTION

In Switzerland, 60% of the land is covered by the Alps with over 1,500 lakes present in these high-altitude regions. Alpine lakes are unique ecosystems with extreme environmental conditions: low nutrient concentrations, low temperatures, ice cover during the wintertime, high UV radiation in summertime, short summertime and drastic light changes between the ice-covered and ice-free season [1]. Despite these harsh conditions, alpine lakes host diverse bacterial communities that play central roles in ecosystem functioning [2]. In aquatic ecosystems, phagotrophic protists and bacteriophages (also known as phages) are major biological factors shaping bacterial populations [3]. However, in alpine lakes bacterial communities are less influenced by protist grazing compared to low-land lakes due to the limited richness and diversity of the latest compared to non-alpine regions [4]. Since bacteriophages are generally highly resistant to extreme environmental conditions, they likely represent the main biological factor shaping bacterial communities in alpine aquatic ecosystems [5].

Bacteriophages are the most abundant microorganisms in aquatic systems [6], [7]. They lead to 20-40% of bacteria mortality and exert a huge selective pressure on their bacteria hosts. Bacteria-viral interactions in this context, are a constant and dynamic back and forth battle [8]. Bacteria adapt their immune systems and develop mechanisms to resist bacteriophage infections, and bacteriophages frequently mutate to counteract bacteria adaptations [9], [10]. Bacteria have developed numerous mechanisms against bacteriophage infections and have clustered them in so-called genomic “defense islands” across their genomes [10], [11]. These defense mechanisms range from the well characterized bacterial CRISPR/Cas system shared by multiple bacterial species to the most recently discovered Bacteriophage Exclusion (BREX) system in *Bacillus subtilis* [12], [13]. They involve phage DNA silencing through small RNA molecules, DNA degradation by restriction enzymes, bacterial programmed cell dead to avoid viral proliferation, and the production of chemical antiviral molecules like viperins [14], [15], [16]. Interestingly, bacterial and human viperins have structural and functional similarities. While bacteria use these molecules to fight bacteriophage infections, humans seem to also use them to fight virus infections [17].

Bacterial antiviral molecules and their mechanisms can be potentially exploited to our advantage against viral pathogens of multiple hosts. Climate change, urbanization and mobility increase the risk of infectious disease emergence [18]. Viral pathogens, in particular, have high mutation rates, large population numbers, short generation times and rapidly become resistant to antiviral drugs and disinfectants [19], [20], [21]. There is a permanent lack of antiviral molecules effective against rapidly evolving and emerging human, animal and plant pathogens [22]. Aquatic ecosystems and, in general, extreme environments are potentially a rich source of microbial secondary metabolites with antimicrobial and antiviral activity. In the early 70s and 80s, Ward *et al*. and Cliver *et al*. described the antiviral activity of fresh water microorganisms and their proteases against Poliovirus, Echovirus-12, Rotavirus SA-11 and Coxsackievirus B5 [23], [24]. More recently, our lab further characterized lakewater bacterial isolates and their proteolytic activity. We found a correlation between viral inactivation and extracellular metalloprotease production [25]. As well, bacterial intra- and extra-cellular polysaccharides have shown antiviral activity against diverse viruses [26]. However, the potential of high-altitude aquatic environments as hotspots for the discovery of new antiviral compounds of bacterial origin remains largely unexplored.

Due to the extreme environmental conditions and the high abundance of bacteriophages in alpine lakes, we hypothesized that bacteria in these environments have developed complex and diverse antiviral mechanisms against bacteriophages that likely contribute to the inactivation of multiple viruses. Thus, we aimed to explore the broad-spectrum viral inactivation by bacteria communities and isolates from Swiss Alpine lakes. We conducted sampling campaigns in 24 alpine lakes located above the tree line in the Swiss Alps. We investigated the inactivation of two human viruses (Influenza A virus and Echovirus 11) in water from selected lakes. In addition, we identified and selected bacteriophage resistant isolates to build two synthetic communities with increased broad-spectrum antiviral activity against the two human viruses and two plant viruses (Potato Virus X and Pepino Mosaic Virus). Our findings highlight high-altitude lakes as unexplored reservoirs of microbial and metabolic diversity. They offer promising opportunities for the discovery of novel broad-spectrum antiviral mechanisms and bioactive compounds, and for the design of synthetic microbial consortia with potential applications in the biocontrol of viral pathogens.

## 2. MATERIALS AND METHODS

### 2.1. Alpine Lake sampling and sample processing

The lakes chosen for this study were high altitude lakes located in the Swiss Alps above 2000 meters of altitude. All samples were collected between September and November 2021. The list of lakes sampled in this project as well as their sampling date and altitude are shown in Table S1. During each sampling campaign, two liters of surface water were collected using plastic containers made of polycarbonate and a telescopic sampling pole. Samples were kept at 4°C until further processing. Once in the laboratory, one liter of lakewater was filtered through 0.45 µm filters (Millipore™) using a conventional vacuum pump to retain protists, bacteria and bacteriophages (associated with the bacteria and organic matter). Water samples with higher turbidity and rich in suspended particles were first pre-filtered through 8.0 µm filters. The 0.45 µm filters were transferred to Bead Tubes of the DNeasy PowerWater kit (QIAgen, Germany). These tubes were frozen at −20°C until nucleic acid extractions were performed using the DNeasy PowerWater kit (QIAgen, Germany) following the manufacturer’s recommendations.

### 2.2. Microbial abundance quantification in alpine lakes

Quantitative polymerase chain reaction (qPCR) assays were used to determine the abundance of picoeukaryotes, bacteria and bacteriophages targeting 18S, 16S and g23 genes, respectively [27], [28]. The 16S gene was targeted to quantify the abundance of bacteria using a Femto Bacterial DNA quantification kit (Zymo Research Corporation) following the manufacturer’s recommendations. The g23 gene encodes for a capsid protein of tailed bacteriophages and was used as a proxy of bacteriophage abundance given that no universal gene is available for viruses[29]. Table S2, shows the sequences of all the primers used in this study.

### 2.3. Metabolic diversity of alpine lake communities

Biolog EcoPlates™ were used to characterize the metabolic profile of bacteria communities found in the studied lakes. Briefly, 1x of M9 minimal media (Sigma-Aldrich) was supplemented with 2mM of MgSO4 (PanReac AppliChem), 0.1 mM of CaCl2 (Thermo Scientific) and 1X of trace metals (Teknova). The content of each well of the EcoPlate™ was resuspended in 100 ul of supplemented M9 minimal. Then, 40 ul of lakewater were added on top of each well. Plates were incubated for 5 days at room temperature in the dark. Optical density of each well was measured at a wavelength of 590 nm using a microplate reader (BioTek – Synergy Mx). Each carbon source is present in triplicate in the EcoPlate™. Average optical density was reported per carbon source for each lake. To calculate metabolic diversity, compounds that yielded an optical density > 0.2 at 590 nm were used to calculate a Shannon index per lake using the following formula:

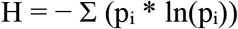

Here, *H* is the Shannon diversity index, *Σ* is the sum, *p*_*i*_ is the proportion of compounds with signal > 0.2 nm and ln is the natural logarithm.

### 2.4. Propagation of human and plant viruses

Echovirus 11 (E11, Gregory strain, ATCC VR737) and Influenza A (IAV) virus were cultured in the laboratory using Buffalo Green Monkey Kidney (BGMK) and Madin-Darby Canine Kidney (MDCK) cells, respectively. Stocks were generated by infecting sub-confluent monolayers of cells, following previously established protocols [30]. For E11 purification, viral particles were harvested from the infected cells through a series of freeze-thaw cycles. Subsequently, to remove cellular debris, the suspensions underwent centrifugation at 3000 × g for 5 minutes. Buffer exchange was performed into PBS using Amicon® Ultra filters in cycles of 4000 × g for 10 minutes until viral suspension was clear. For IAV purification, culture supernatant from the infected cells was cleared by centrifugation (2,500 × g, 10 min). Virus was concentrated and purified by pelleting through a 30% sucrose cushion at 112’400 × g in a SW31Ti rotor (Beckman) for 90 min at 4°C. The resulting pellets were soaked overnight at 4 °C in PBS (ThermoFisher, 18912014) and then fully resuspended by pipetting up and down. Pepino Mosaic Virus (PepMV) and Potato Virus X (PVX) were obtained using tobacco plants (*Nicotiana benthamiana* and *Nicotiana tabacum Xanthi*) mechanically inoculated with an isolate of PepMV in an Agroscope greenhouse in Changins (Nyon, Switzerland). For virus propagation, symptomatic leaves were harvested, homogenized in phosphate buffer, and clarified by low-speed centrifugation. Virus abundance in the obtained stocks was confirmed by electronic microscopy. All virus suspensions were aliquoted and stored at −80 °C until further use.

### 2.5. Enumeration and quantification of human viruses

The concentration of infectious viruses was determined using a most probable number (MPN) infectivity assay for E11 and plaque assays for IAV. For MPN, 5 replicates per sample were used as described previously [20]. Plaque assays were performed using MDCK cells as described previously [31].

For quantification of human viruses, qPCR targeting Influenza A Matrix or Echovirus 11 viral protein 1 were performed (Table S2). For both viruses, RNA extractions from samples were performed, using the QIAamp Viral RNA Mini extraction kit (52906, QIAGEN GmbH, Hilden, NW, Germany) according to manufacturer’s instructions. For E11 quantification, RT-qPCR was performed on a Mic qPCR Cycler (Bio Molecular Systems) using the One Step SYBR® PrimeScript™ RT-PCR Kit (Takara). Each reaction (20 μL) contained 3 μL of template, 10 μL of 2× One Step SYBR RT-PCR Buffer III, 0.4 μL of TaKaRa Ex Taq HS (5 U μL−1), 0.4 μL of PrimeScript RT enzyme mix II, 250 nM of each primer and 5.2 μL of water. The thermocycling conditions were as follows: RT at 42 °C for 5 min, 10 s at 95 °C, followed by 40 cycles of 95 °C for 5 s, 52 °C for 20 s and 60 °C for 30 s. Each sample was run once, and the Cq values were determined using the micPCR software (v2.10.0; Bio Molecular Systems)[32]. For IAV quantification, extracted nucleic acids were eluted in elution buffer, and stored at −20°C until analysis. Reverse-transcription and amplification were performed using the One Step PrimeScript™ RT-PCR Kit (RR064A, Takara Bio Inc., Kusatsu, JP-25, Japan) using primers targeting the Influenza A Matrix gene (Table S2). The RT-qPCR mix was composed of 7.5 μL of 2x One-Step SYBR RT-PCR Buffer, 0.3 μL of Takara Ex Taq HS (5 U/μL stock), 0.3 μL PrimeScript RT enzyme mix, 0.3 μL forward and reverse primers (10 μM stocks), and 3.3 μL RNase-free water, to which 3 μL of extracted RNA sample was added. The RT-qPCR was performed in a Mic Real-Time PCR System (Bio MolecularSystems, Upper Coomera, QLD, Australia) with the following parameters: 2 min at 50 °C, 10 min at 95 °C, 15 seconds at 95 °C followed by 60 sec at 60 °C for 40 cycles. A final step from 55 °C to 95 °C at 0.3 °C/s was performed for the melting curve analysis. A Gblock gene fragment (Gblock SA, Ath, WHT, Belgium), as described in Olive et al. (2022), was used to create a standard curve for quantification over a range of 10 to 10^7^ genomic copies per microliter (GC/μl)[33].

### 2.6. Inactivation of human viruses in lakewater

Inactivation experiments were conducted in lakewater filtered through 0.8 µm (Millipore™) (to remove heterotrophic protists) as described above. Specifically, 50 ml of filtered water were spiked with 100 µl of IAV and E11 to the same initial concentration (10^5^ genome copies per ml). Then, 5 ml of spiked water was placed in 6-well plates in three replicates. The plates were incubated for 7 days at room temperature and 500 µl aliquots were taken on days 0, 3 and 7. These aliquots were frozen at −20°C until analyzed. Viral nucleic acids were extracted using the QIAamp Viral RNA Mini extraction kit (52906, QIAGEN GmbH, Hilden, NW, Germany) and virus concentrations were determined by qPCR throughout the experiment or by cell culturing as described above.

### 2.7. Development of synthetic bacteria communities with antiviral activity

To develop synthetic bacteria communities with enhanced antiviral activity, we first aimed to identify specific bacteria species with antiviral properties in the lake water samples. To this end, we built a culture collection of isolates from all the sampled lakes. One liter of water from each lake was 0.45 *μ*m filtered. Filters were transferred to LB agar plates and incubated at room temperature for 72 hours. Then, single colonies were transferred to new LB plates. Serial transfers of isolated colonies into sterile LB plates were conducted until pure cultures were obtained. A total number of 223 pure bacterial isolates were obtained using this methodology. LB media was purchased from Invitrogen. Bacteria isolates were identified performing 16S rRNA gene sequencing using the primers 27 F and 786 R (Table S2). Samples were sent for conventional Sanger sequencing to Fasteris (Genesupport SA, Geneva, Switzerland).

To identify which of the obtained isolates showed antiviral activity, we conducted phage susceptibility tests using the bacteriophage fraction obtained from each studied lake. Specifically, phage enrichments were conducted for each bacteria isolate obtained in this study (n=223) following state-of-the-art protocols [34], [35]. The phage cocktail used to infect each bacteria isolate was a 1:1 mixture of all the control waters obtained from each lake which is called “Phage Mix” hereafter. This process was done to increase the chances for at least one phage type in the mix to be capable of infecting each bacteria isolate. To assess bacterial susceptibility to the “Phage Mix”, fresh cultures of each bacteria isolate were grown overnight in 5 ml of LB. Once bacteria cultures had an optical density at 600 nm around 0.6-0.8 (exponential phase), they were spiked with 1 ml of the phage mix. The tubes were then incubated at room temperature for 15 min and then gently agitated (50 rpm) for 2 more hours. Then, chloroform was added to the bacteria-phage co-cultures (hereafter named “Phage enrichments”) which were centrifuged for 10 minutes at 5000 rpm and filtered at 0.2 µm to remove all bacteria. This process was done for each bacteria isolate.

To compare phage susceptibility of bacteria isolates, the phage titer yield in the “Phage Mix” and in each enrichment was determined by qPCR targeting the g23 gene [34], [35]. Bacteria cultures with phage concentrations equal or higher compared to the phage mix were considered “susceptible” and retained to build the control communities, while those yielding inferior concentrations were considered “resistant” and retained for the design of two different synthetic communities with enhanced antiviral properties. This approach assumes that resistant bacteria limit phage replication, so lower phage titers in the enrichment experiments reflect the bacteria’s ability to inhibit phage proliferation.

### 2.8. Alpine synthetic communities against human and plant viruses

To confirm the enhanced antiviral properties of the obtained synthetic communities, three groups of 25 bacterial isolates (2 synthetic communities with enhanced activity and one control group) were mixed and used to assess their inactivation activity against E11 and IAV. Before the inactivation assays, isolates were independently grown in LB overnight at room temperature to avoid competition and were mixed in equal parts in a final volume of 50 ml. Once the bacteria isolates were mixed together, inactivation experiments against E11 and IAV were conducted as previously described. Two bacterial pools were composed of 25 unique bacterial isolates with decreased susceptibility to the “Phage Mix” and one control was composed of 25 bacteria isolates with increased phage susceptibility. Specifically, 50 ml of mixed bacteria isolates were diluted 100x using sterile lakewater and spiked with 100 µl of IAV and E11 to the same initial concentration (10^5^ genome copies per ml). Then, 5 ml of spiked media with the isolates was placed in 6-well plates in five replicates. The plates were incubated for 4 days at room temperature. For inactivation assays against plant viruses, the most inactivating community against human viruses was selected. All the community members were grown as previously described and mixed the day of the assay. Previously purified PepMV and PVX viruses were independently spiked into the bacteria solution and incubated at room temperature for four days. After incubation, the viral-bacterial solutions were used to infect 75 tobacco plants. Inoculated plants were maintained under controlled greenhouse conditions until infections symptoms appeared (approx. 15 days). A group of uninoculated plants were placed in the same green house as plant controls. In addition, a group of 16 plants were inoculated only with the bacterial pool as a control of bacteria pathogenesis. Once symptoms appeared, two leaves were harvested per plant and used to confirm the presence of the viruses by ELISA.

### 2.9. 16S amplicon sequencing of alpine synthetic communities

Once bacterial isolates were mixed, a sample from each community were taken for DNA extraction and 16S amplicon sequencing. Sample was 0.22 µm filtered and DNA from filters was extracted using DNeasy PowerWater Kit (Qiagen, Hilden, Germany) as previously described. DNA extracts and blanks were then used for 16S rRNA gene amplification using the 27F and 1492R (Table S2). The PCR reaction was carried out in a 50-μL volume containing 20 ng of input DNA and 25 μL of LongAmp Taq 2× master mix (New England Biolabs, Ipswich, MS, USA). Each PCR product was purified using Agencourt AMPure XP beads (Beckman Coulter, Brea, CA, USA) and quantified using Qubit™ (dsDNA quantification, Broad Range). The sequencing library was prepared using the Native Barcoding Kit SQK-NBD114.24 (Oxford Nanopore Technologies, Oxford, UK) following the manufacturers’ instructions. The library was loaded into a FLO-MIN106 flow cell (Oxford Nanopore Technologies) and sequenced using a MinION Mk1C sequencing device for 72 hours. Sequencing files were basecalled using Dorado (v0.9.1) basecaller. FASTQ files were input into the MetONTIIME workflow [36] to estimate bacterial relative abundance and assign taxonomy. Feature tables and count matrix were imported into R and analysed using the *microeco* package [37].

### 2.10. Data analysis and visualization

Analysis of all the data obtained during the project was done with the *dyplr* and *tidyverse* packages RStudio [38]. All the plots were obtained in Rstudio using the *ggplot* package [39]. Correlation matrices and statistical analysis were performed using the Pearson correlation coefficient function in RStudio.

## 3. RESULTS AND DISCUSSION

### 3.1. Microbial abundances and metabolic diversity in Swiss alpine lakes

A total of 24 alpine lakes were sampled in the Swiss Alps from eight distinct regions in the canton of Valais and one region in the Bernese Oberland (Figure 1A). The abundance of bacteria, eukaryotes and tailed bacteriophages was determined in each lake to identify those showing a higher ratio of bacteriophages to bacteria and eukaryotes. We found great variability of microbial abundances between samples (Figure 1B). Bacteria concentrations (16S) ranged from 300 genome copies/liter (gc/l) in Lac Bleu (Arolla) to more than 10^8^ gc/l in Spilsee (Bellwald). Eukaryotic DNA (18S) ranged from 100 gc/l in Lac Bleu to 10^9^ gc/l in Stellisee and Leisee (Zermatt). The concentration of tailed bacteriophages (g23) varied between < 10 gc/l in Small Lac de Vaux (Verbier) to 10^7^ gc/l in Lengsee (Bellwald). We calculated the ratio of tailed bacteriophages over the sum of bacteria and eukaryotes concentration (measured in gc/l) in each lake, shown in Figure 1C. At the time of sampling, few of the lakes had particularly high ratios of tailed bacteriophages per bacteria and protists. Lakes Mossjesee, Guggisee, Marjelensee and Salanfe had ratios of 0,12, 0,15, 0,10 and 0,69, respectively, compared to an average ratio of 0,01 in the other 20 lakes. The ratio of tailed bacteriophages to bacteria and eukaryotes was 60-times higher in Lake Salanfe compared to the average ratio of the rest of the lakes. In agreement with earlier studies, the variability in microbial abundances among the studied lakes may reflect local environmental (e.g. ionic concentrations, nutrients, temperature and pH) and biological influences. Identifying and characterizing the influence of these specific factors was beyond the scope of this work [40].

**Figure 1.**
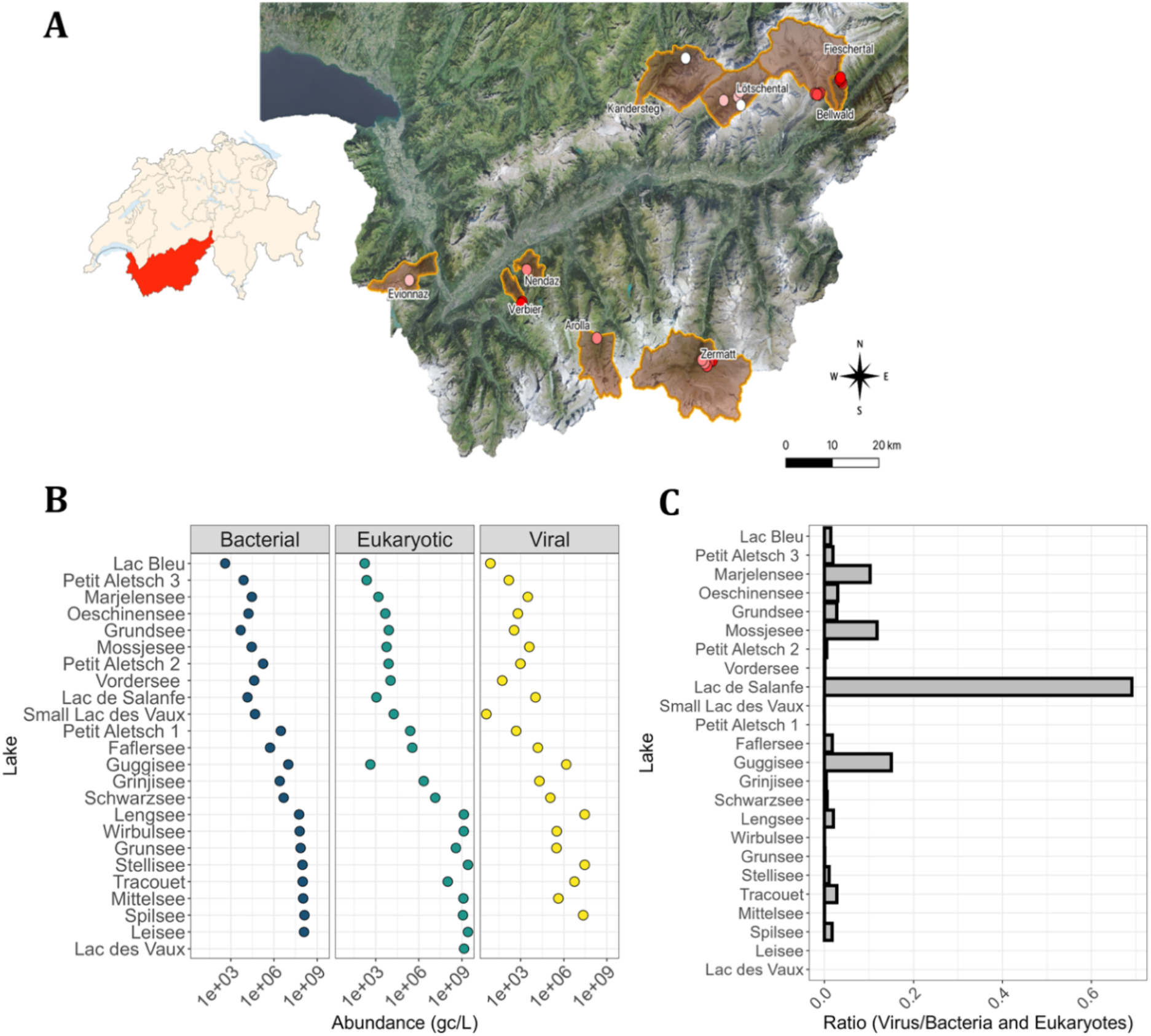
Microbial abundance of 24 Swiss Alpine Lakes. **(A)** In total 24 lakes located above 2000 meters of altitude were sampled from August to November 2021. **(B)** Total DNA was extracted from lakewater samples and the genome copies per litre (gc/L) of bacteria bacteria, eukaryotes and tailed phages was determined by quantitative PCR (qPCR) using primers against 16S, 18S and g23 genes, respectively. **(C)** The ratio of bacteriophages to the sum of bacteria and eukaryotes were calculated per lake using the abundance (gc/L) of g23 over the sum of abundance of 18S and 16S genes.

To characterize functional traits of the microbial communities in the sampled lakes that could provide insights into their antiviral activity, we assessed carbon metabolism using Biolog EcoPlates. These plates contain 31 compounds classified as carbohydrates, amino acids, carboxylic acids, amines or alcohols (Figure 2). Based on the number of metabolized carbon sources, we calculated Shannon diversity indexes to determine the metabolic diversities per each lake (Table S3). Lakes Bleu, Petit Aletsch 1 and Schwarzsee had the highest metabolic diversity (Shannon indexes > 3). Microbial communities present in these three lakes metabolized substrates in all six categories (carbohydrates, amino acids, carboxylic acids, amines and alcohols). In contrast, lakes Stellisee and Leisee were the least metabolically diverse (Shannon indexes < 1), as microbial communities from these lakes were unable to metabolize most of the substrates present.

**Figure 2.**
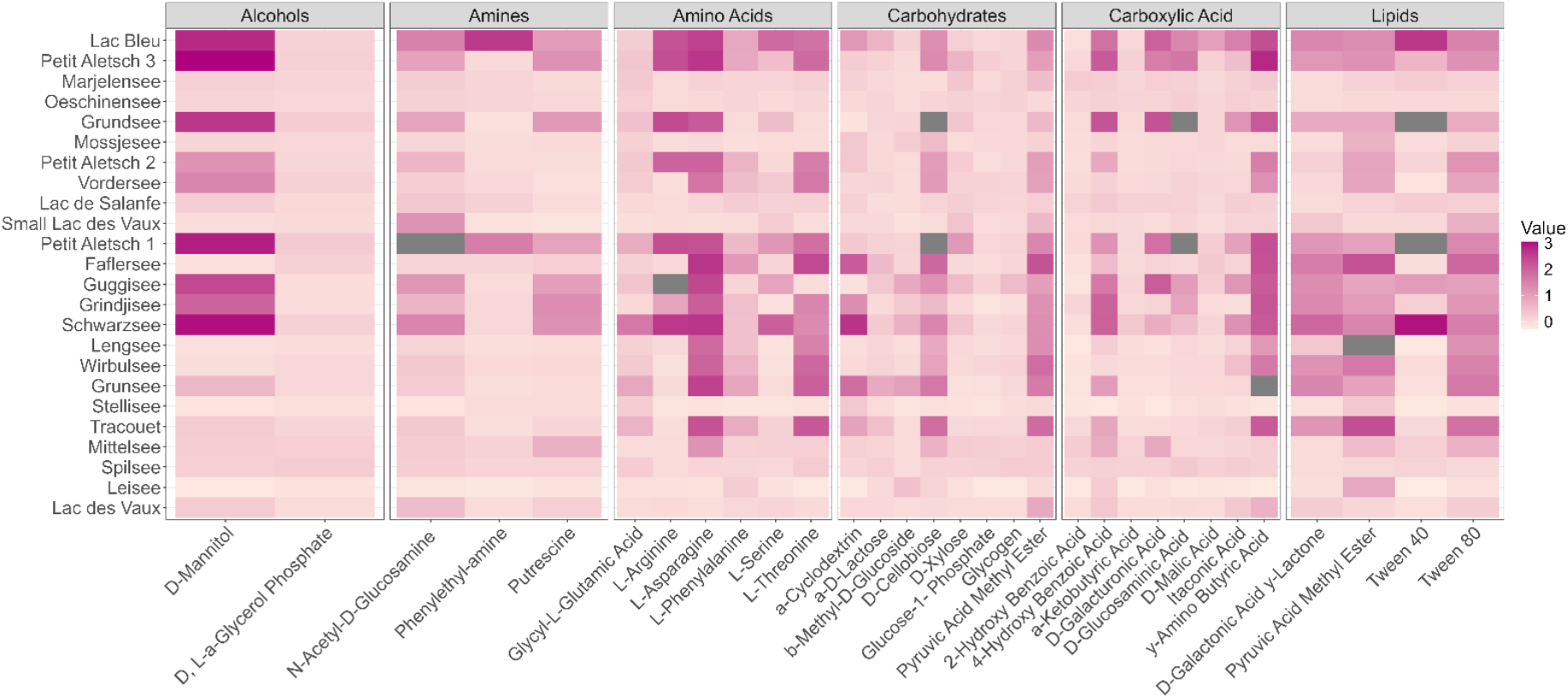
Metabolic profiling of microbial communities in Swiss Alpine Lakes. **Water** from each lake was used to assess the capacity of each lake’s microbial community to degrade the 31 different substrates present in Ecolog plates. Each lakewater was inoculated into an Ecolog plate and incubated for five days at room temperature. Optical density was measured at 590 nm and averaged for each substrate from three replicates (n=3).

### 3.2. Microbial communities in alpine lakes show antiviral activity against human viruses

We have previously reported that bacterial communities from Lake Geneva inactivate human enteric viruses [25]. To confirm whether bacteria in Swiss alpine lakes also show antiviral activity against human viruses, we conducted inactivation experiments using two structurally different viruses: echovirus 11 (E11) and influenza A virus (IAV). For these assays, we selected four lakes with high proportion of tailed bacteriophages per bacteria and eukaryotes (Guggisee, Mossjesee, Marjelensee and Salanfe), and four lakes with low proportion (Lac Bleu, Schwartzsee, Small Lac de Vaux and Mittelsee) to evaluate whether elevated phage-to-host ratios are indicative of increased antiviral activity. We focused on the antiviral activity of bacteria present in the water. Therefore, the water used for these assays was filtered through a 0.8 µm membrane to remove protists, also known to inactivate human viruses [30].

We observed higher E11 inactivation compared to IAV in all the tested lakes (Figure 3). E11 was inactivated > 1 log10 in five out of eight of the lakes (Figure 3A) while IAV was inactivated around 1 log10 in all lakes (Figure 3B). These findings are consistent with previous observations on the inactivation of human pathogens in freshwater [30], [41]. Out of the eight lakes, lake Mossjesee, an artificial reservoir resulting from glacial meltwater, had the greatest inactivation against both viruses (2.17 average log10 decay for E11; 1.03 log10 decay for IAV) over 7 days of incubation. E11 inactivation in water from this lake was significantly higher compared to lakes Salanfe and Mittelsee (P = 0.0387 compared to Salanfe and P = 0.0199 compared to Mittelsee). We did not observe a correlation between viral inactivation and the proportions of bacteriophages per bacteria and eukaryotes. However, two out of four of the chosen lakes with increased ratios of tailed bacteriophages (Mossjeesee and Marjelensee) showed the greatest inactivation against E11. Similarly, we did not identify any correlation between antiviral activity and lake metabolic diversities (Figure S1). Further research investigating ecosystem parameters associated with higher viral inactivation in alpine lakes is necessary.

**Figure 3.**
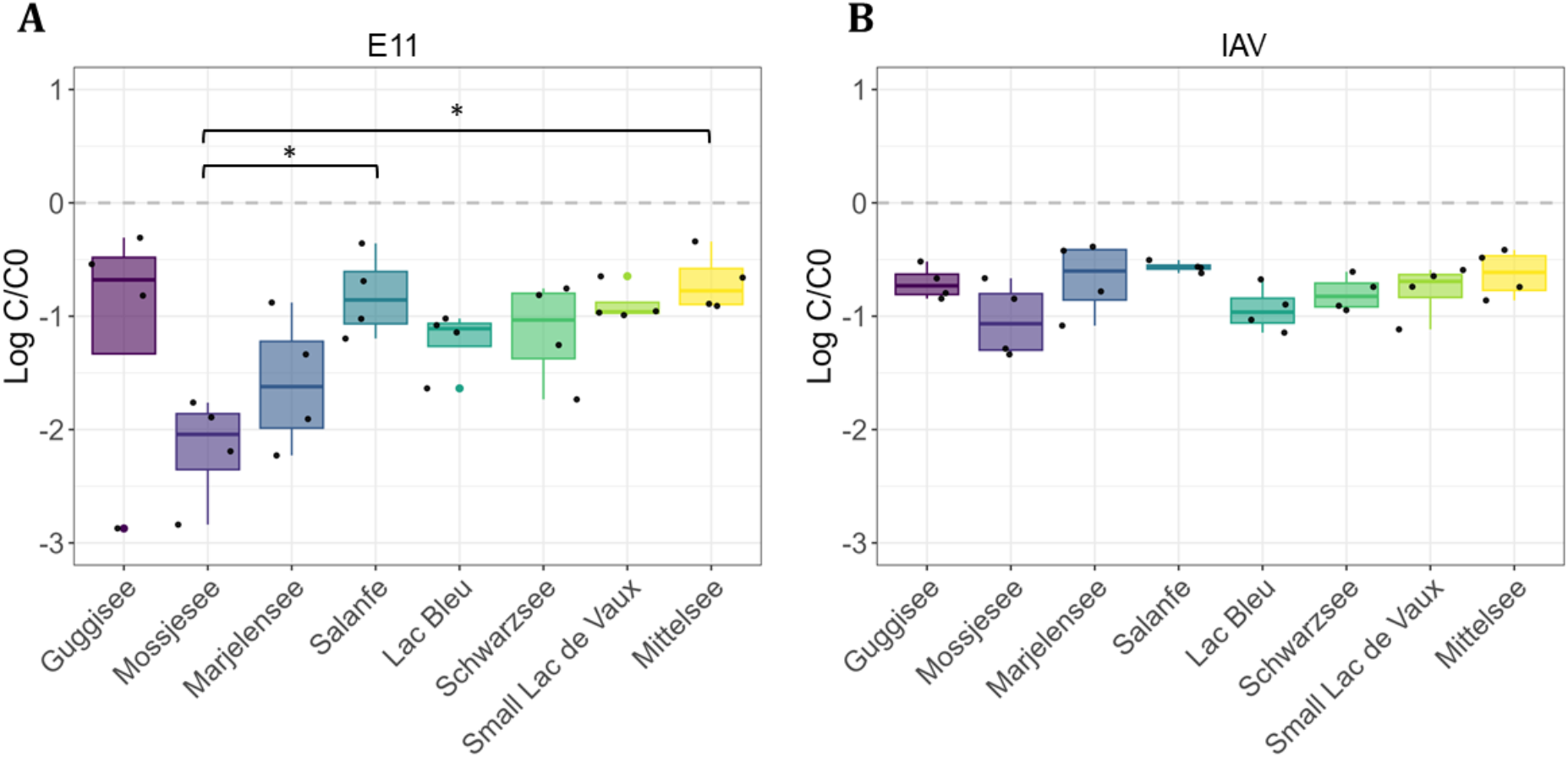
Inactivation of Human viruses in selected Alpine Lakes. **(A)** Echovirus 11 (E11) or **(B)** Influenza A virus (IAV) was inoculated into 0.8 μm filtered water from eight Alpine lakes and incubated for seven days at room temperature. Viral inactivation was determined by comparing the viral genome copies per ml (gc/ml) on day seven to day zero (Log C/C0) using quantitative PCR (qPCR). Each viral inactivation assay was performed in five replicates (n = 5). A one-way ANOVA with Tukey honest significant difference in R Studio was used for statistical analysis between viral decay values. Significant differences are indicated as follows: ***P < 0.001, **P < 0.01, and *P < 0.05.

### 3.3. Alpine lakes host bacteria with low susceptibility to bacteriophage infections

To identify and characterize specific bacteria with potential antiviral activity, we isolated single bacteria species from each lake in LB agar (Figure 4A), regardless of the phage-to-host ratios. In total, we obtained 223 different bacteria isolates across the 24 alpine lakes (Figure 4B). The number of bacterial isolates obtained varied per lake, ranging from 4 to 19. Lakes Guggisee and Oeschinensee yielded the highest number of isolates (19 and 16 cultures respectively), together accounting for approximately 15% of the total isolates. Lakes Stellisee and Wirbulsee yielded the lowest numbers of bacterial isolates (only 4 isolates from each of both lakes). Interestingly, Lakes Guggisee and Oeschinensee showed the lowest bacterial abundance (Figure 1B), while lakes Stellisee and Wirbulsee were within the lakes with the greatest bacterial abundance.

**Figure 4.**
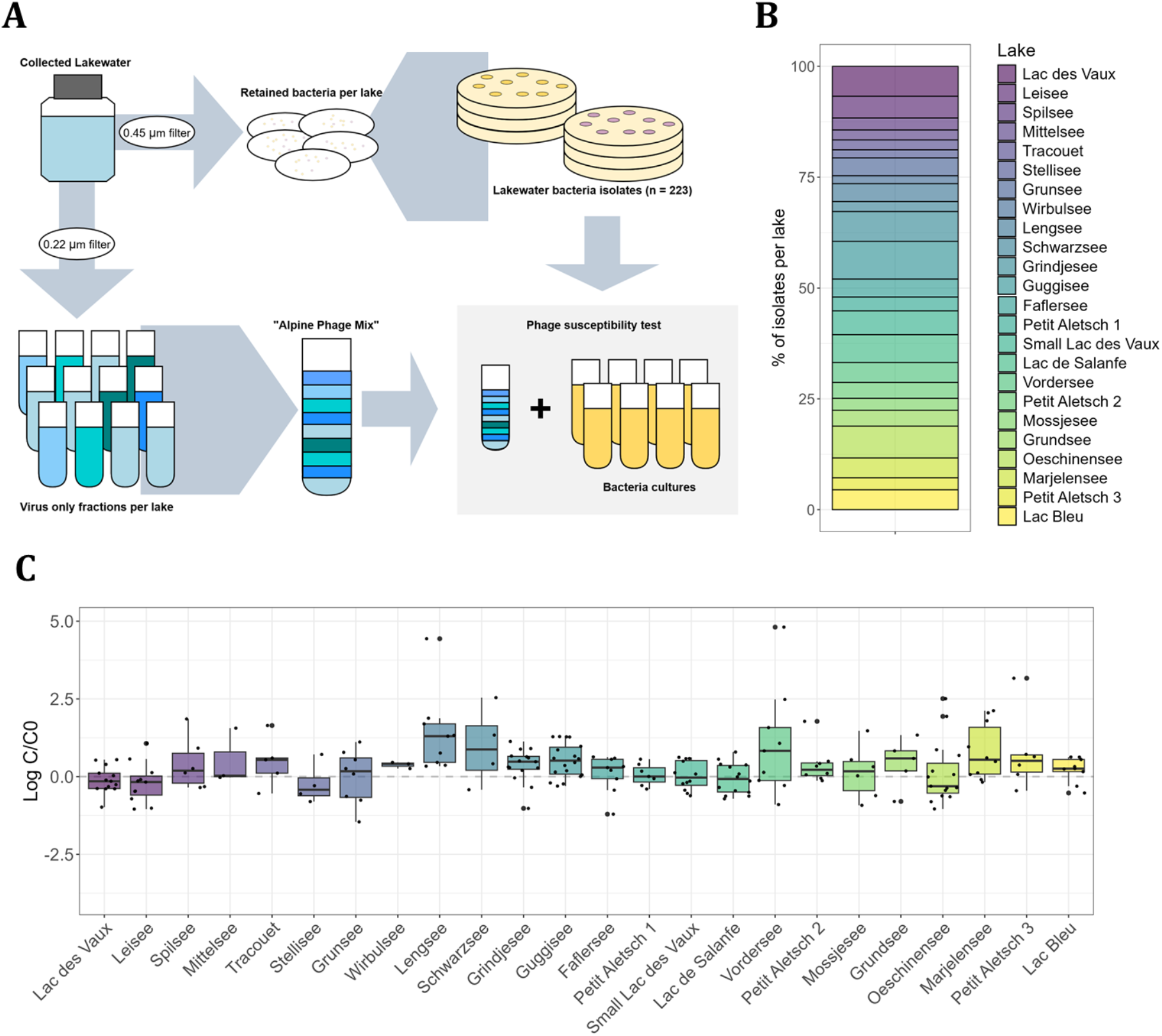
Phage susceptibility of Alpine bacterial isolates. **(A)** An “Alpine Phage mix” was generated by mixing lakewater samples from each lake. In parallel, to isolate bacteria from each lake, one litre of lakewater was 0.45 mm filtered. Filters were transferred to LB agar plates and incubated at room temperature for 72 hours. **(B)** A total of 223 bacterial isolates were recovered and identified using 16S gene sequencing. The number of bacterial isolates obtained varied per lake. **(C)** To assess bacterial susceptibility to the Alpine Phage mix, each bacterial isolate was grown in LB and spiked with one ml of the mix. After five hours, bacterial cultures were recovered, spun down and filtered. DNA was extracted at 0 and 5 hours and qPCR against the g23 gene was used to compare phage titers between these two time points (Log C/C0).

We hypothesized that bacterial isolates with lower susceptibility to bacteriophage infections would show increased antiviral activity against human and plant viruses. To identify bacteria with decreased susceptibility to bacteriophage infections and test our hypothesis, we created a “Phage Mix” by pooling virus fractions (0.22 µm filtered lakewater) from the surface water samples collected in the 24 lakes. This generated a phage cocktail that included a large abundance of diverse bacteriophages potentially capable of infecting most of the bacteria isolates (Figure 4A). All 223 isolates were grown independently and spiked with the alpine “Phage Mix”. After co-incubating the bacteria isolates and the phage mix, we compared the final concentration of bacteriophages to the initial concentration in the inoculum (Figure 4C).

Most of the isolates (148 specifically) showed susceptibility to the Phage Mix as indicated by an increase in the bacteriophages’ titer after the incubation period. Conversely, we identified 75 isolates with decreased susceptibility indicated by a decrease in the concentration of bacteriophages after co-incubations. Lake Oeschinsee (9 isolates), Lac de Vaux (8 isolates), Leisee (8 isolates) and Lac de Salanfe (8 isolates) had the greatest number of phage resistant isolates. Interestingly, these lakes had the lowest metabolic diversity (Shannon 0.98 to 1.69) compared to other lakes. In terms of bacteriophage to bacteria and eukaryotes ratios, we did not observe any relation between bacteriophage resistant isolates and the calculated ratios. However, we noticed that Lac de Salanfe had the highest calculated ratios amongst all sampled lakes (Figure 1C).

Our bacteriophage susceptibility tests on alpine bacteria narrowed the initial set of 223 isolates to 75 potentially bacteriophage resistant strains. It is important to take into account that some factors and choices made such as: media selection for bacterial isolation; possible absence of phages capable of infecting certain isolates in the Phage Mix [7][42]; and using g23 qPCR as an indicator of total phage concentration may have influenced our results. However, the identified bacterial subset represented promising candidates for further characterization and was used to develop synthetic communities with enhanced viral inactivation activity.

### 3.4. Phage-resistant synthetic communities inactivate human and plant viruses

Previous studies performed in our laboratory showed greater viral inactivation in lakewater samples compared to single bacteria cultures, and a correlation between increased microbial diversity and enhanced inactivation [25], [43]. Therefore, we aimed to design microbial synthetic communities using isolates classified as “bacteriophage-resistant” and assess their antiviral activity against viral pathogens with different structures and hosts. We created two synthetic communities of 25 members each (communities 1 and 3 – Figure 5). In parallel, we created a control community using 25 bacteriophage-susceptible bacteria (Control community). The three communities were spiked with either E11 or IAV and incubated for 4 days. After co-incubation, we confirmed that the synthetic communities of bacteriophage-resistant bacteria (communities 1 and 3) had enhanced antiviral activity compared to the bacteriophage-susceptible (control) community (Figure 5A). In both cases, we observed significantly (P value < 0.01) more inactivation between the communities (1 and 3) and the control but no significant difference between communities 1 and 3. In addition, we observed that E11 had greater inactivation compared to IAV (E11 ~ 3 log10; IAV ~ 1 log10, in four days). Noteworthy, the synthetic communities showed greater inactivation against both human viruses as compared to the inactivation we previously observed in lakewater (~ 2 log10 in seven days).

**Figure 5.**
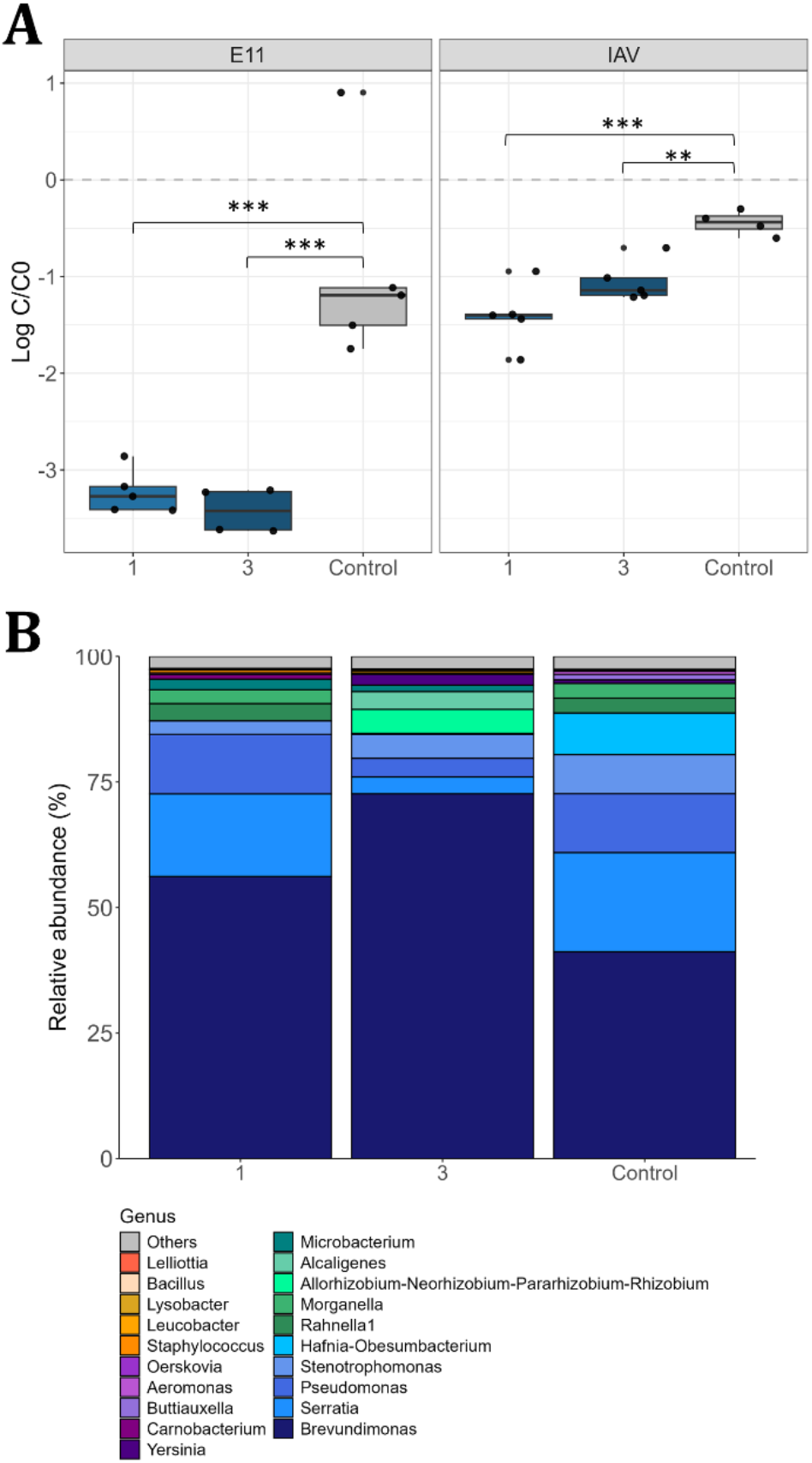
Alpine synthetic communities inactivate human virus. Bacteria isolates were mixed to generate synthetic communities of 25 members. Communities 1 and 3 were created by mixing lakewater bacteria with less susceptibility to the “Alpine Phage Mix”. A control community was generated by mixing bacteria with the least susceptibility out of the 223 isolates. **(A)** To assess the inactivation capacity of each bacterial community, Echovirus 11 (E11) and Influenza A virus (IAV) were spiked into each community. Viral inactivation was determined (Log C/C0) using quantitative PCR (qPCR) as previously described after 4 days. Each viral inactivation assay was performed in five replicates (n = 5). **(B)** Community composition at the end of the experiment was determined by extracting DNA from each community and sequencing the 16S gene using Oxford Nanopore Technologies (ONT). A one-way ANOVA with Tukey honest significant difference in R Studio was used for statistical analysis between viral decay values. Significant differences are indicated as follows: ***P < 0.001, **P < 0.01, and *P < 0.05.

We determined the bacteria composition of the three communities (1, 3 and control) before viral inoculation using 16S amplicon sequencing and confirmed the identity of individual bacteria using Sanger sequencing (Figure 5B). We found high relative abundance of genera *Brevundimonas, Serratia, Pseudomonas* and *Stenotrophomonas* species across the three communities (control and communities 1 and 3). On the other hand, genera *Alcaligenes* and *Microbacterium* species were shared between communities 1 and 3 but were not present in the control community. We also identified bacteria uniquely found in each community. *Bacillus* sp. and *Staphylococcus* sp. were uniquely present in community 1, and *Allorrhizobium* species were uniquely found and abundant in community 3. Some of these bacteria genera have previously shown antiviral activity against plant pathogens. For example, a protease purified from a strain of Serratia marcescens protected tobacco plants against Tobacco Mosaic Virus (TMV) [44]. In addition, secondary metabolites from *Bacillus subtilis* also have proven antiviral activity against RNA viruses and a pseudorabies virus (PVR) [45], [46]. These antiviral molecules have been previously described in single strains but their production and role in inactivation in microbial communities remains unknown.

Lastly, to identify some mechanisms behind the observed antiviral activity, we assessed protease secretion by individual bacteria from each community. Bacteria isolates were individually grown in skim milk agar (Figure S2A) and proteolytic activity was determined by measuring degradation halos around individual colonies. We did not observe a difference between the proteolytic activity of the community members present on each of the phage resistant and susceptible communities (Figure S2B). Viral degradation was first described by Cliver et al. (1972) and Ward et al. (1986). They showed antiviral activity of bacteria against enteroviruses and concluded that viral RNA was easily degraded after cleavage of viral proteins by proteolytic bacterial enzymes [24], [47]. Other studies on the fish pathogen Koi-Herpesvirus have shown significantly faster inactivation with bacteria present compared to bacteria free samples [48], [49]. They also suggest that proteases were likely responsible for the observed inactivation [48]. We were unable to find a correlation between viral inactivation and the proteolytic activity of the community members. This indicates that either the assays used may have not captured the specific proteolytic enzymes responsible for viral inactivation or other bacterial mechanisms also contribute to the observed antiviral activity. Further studies are needed to elucidate the specific mechanisms underlying viral inactivation by the enhanced synthetic communities generated in this study, as well as the contribution of each individual species in the community.

Since our previous results indicated that the developed synthetic communities exhibit antiviral activity against bacteriophages and human viruses, we hypothesized that they may possess broad-spectrum antiviral properties. To test this hypothesis, we performed viral inactivation assays against Potato Virus X (PVX) and Pepino Mosaic Virus (PepMV) using synthetic community 3 (Figure 6). For these assays, the synthetic community was spiked with either plant virus, and then incubated at room temperature for four days. In parallel, both viruses were also diluted in PBS and incubated under the same conditions. These suspensions were used to mechanically inoculate tobacco plants. After 15 days of incubation, we found a 72% reduction in viral infectivity of PVX incubated with community 3 compared to PVX alone (Figure 6A and B). In contrast, we did not observe viral reduction of PepMV incubated with the bacterial community (Figure 6 C and D). These results suggest that community 3 exhibits broad-spectrum antiviral activity but also displays some specificity in the extent of viral inactivation across different viruses. As controls, we included 10 untreated plants to evaluate crossed contamination in the greenhouse (Plant controls - Figure 6E). We also included 16 plants inoculated with community 3 to assess bacteria pathogenesis against plants (Figure 6F). One of our plant controls was positive for PVX and two of the plants inoculated with the bacteria community showed signs of infection. These results confirmed our working hypothesis and provide for the first time a link between resistance against phage infections and viral inactivation.

**Figure 6.**
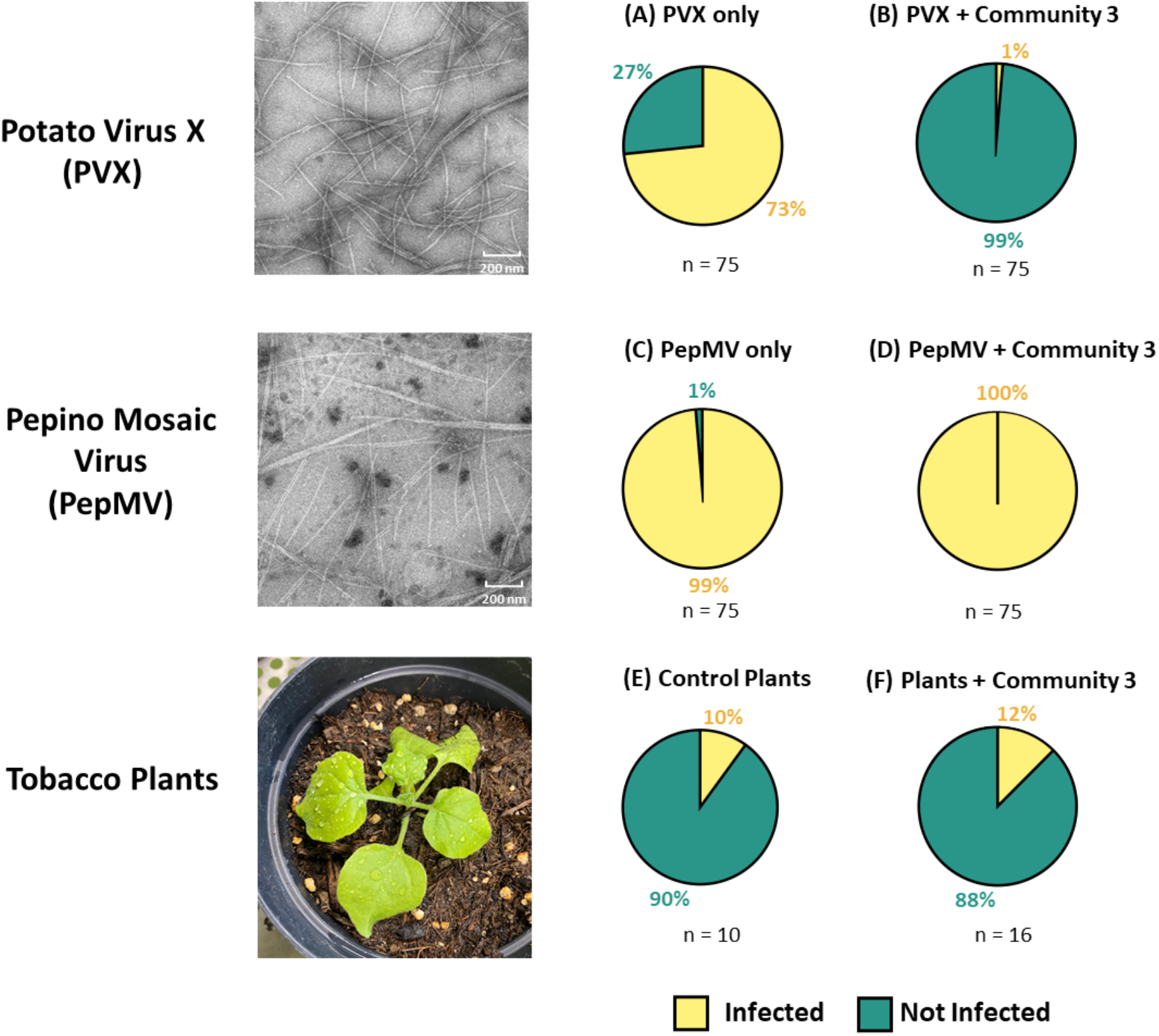
Bacterial Pool 3 shows antiviral activity against Potato Virus X (PVX). The antiviral activity of “Pool 3” was assessed against Potato Virus X (PVX) and Pepino Mosaic Virus (PepMV). Both viruses were purified from tobacco plants and diluted into a PBS solution (A and C) or the “Pool 3” bacterial community (B and D). Virus only and Virus + Pool 3 were incubated at room temperature for 4 days and used to infect tobacco plants. After 15 days, leaves were taken from each plant to perform an ELISA assay against both viruses (in duplicate). Plant controls were incubated and processed as treated plants (E). Some plants were inoculated with the bacteria mix to assess bacteria pathogenicity against plants (F).

### 3.5. Perspectives

In this study, we developed and applied an innovative strategy to construct two synthetic bacterial communities with enhanced broad-spectrum antiviral activity from environmental samples collected in 24 Swiss alpine lakes. We further demonstrated that alpine lakes exposed to extreme environmental conditions harbor bacterial populations with antiviral properties. This was consistent with our previous observations in lowland lakes such as Lake Geneva. Notably, our results provide the first evidence linking resistance to indigenous environmental bacteriophages with inactivation of viral pathogens infecting diverse hosts, highlighting potential avenues for future biotechnological applications. Future research will aim to elucidate the molecular mechanisms underlying the broad-spectrum viral inactivation reported here.

## FUNDING AND ACKNOWLEDGEMENTS

This work was supported by the project ANTIVIRALPS (“Screening for novel antiviral compounds in alpine and polar lakes”) funded by the EPFL Science Seed Fund 2021, and the project HYDROVIR (“Optimizing the use of alpine lake water bacteria with antiviral properties to combat viral diseases in hydroponic cultures”) funded by the EPFL ENABLE Grant, 2022. The authors gratefully acknowledge Nathalie Dubuis for her assistance with plant virus preparation and sample analysis in Agroscope.

---

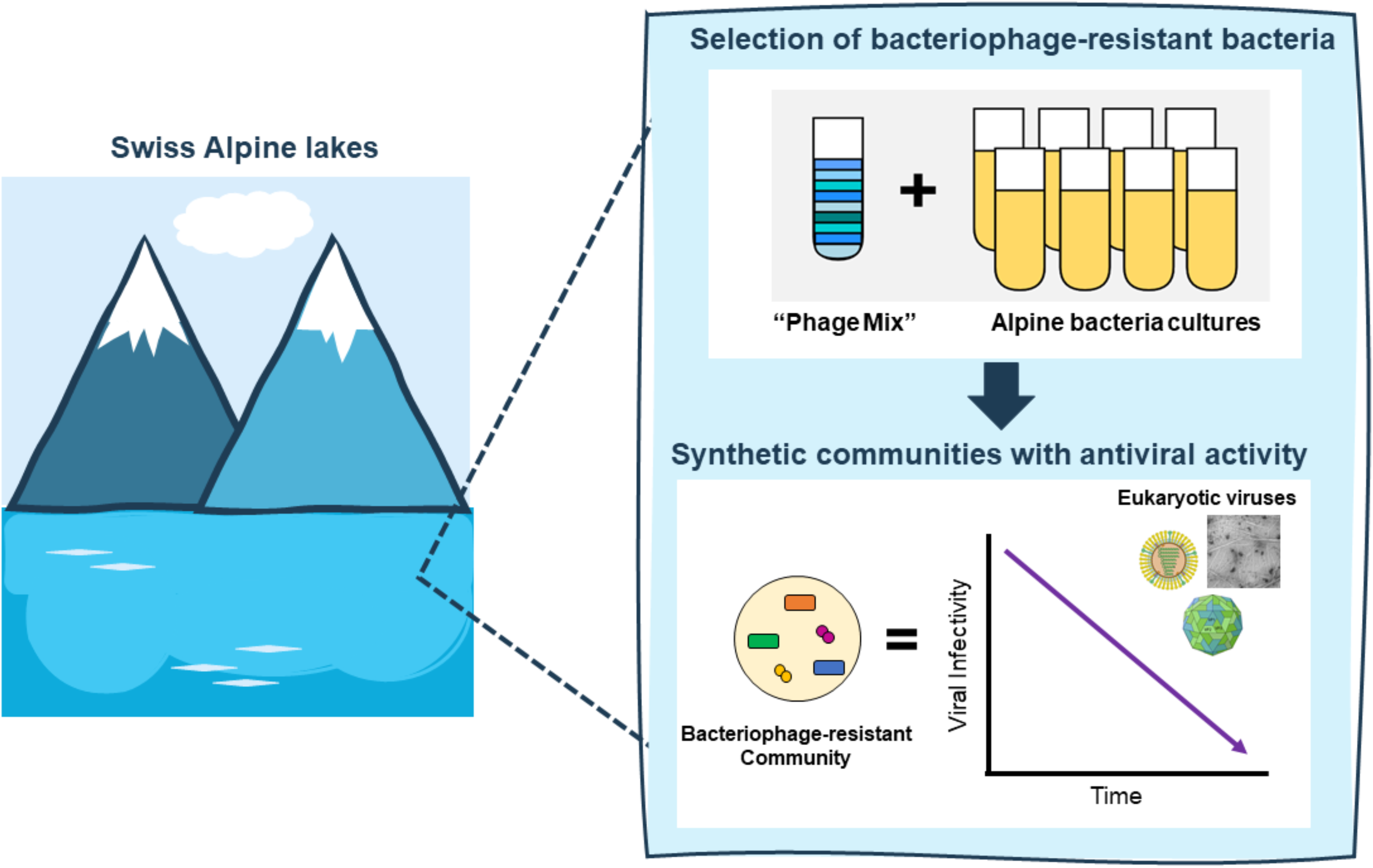

