## Supplemental_information for "Harnessing Alpine Lake Bacteria for the Development of Synthetic Communities with Broad-Spectrum Antiviral Activity"

### Figures

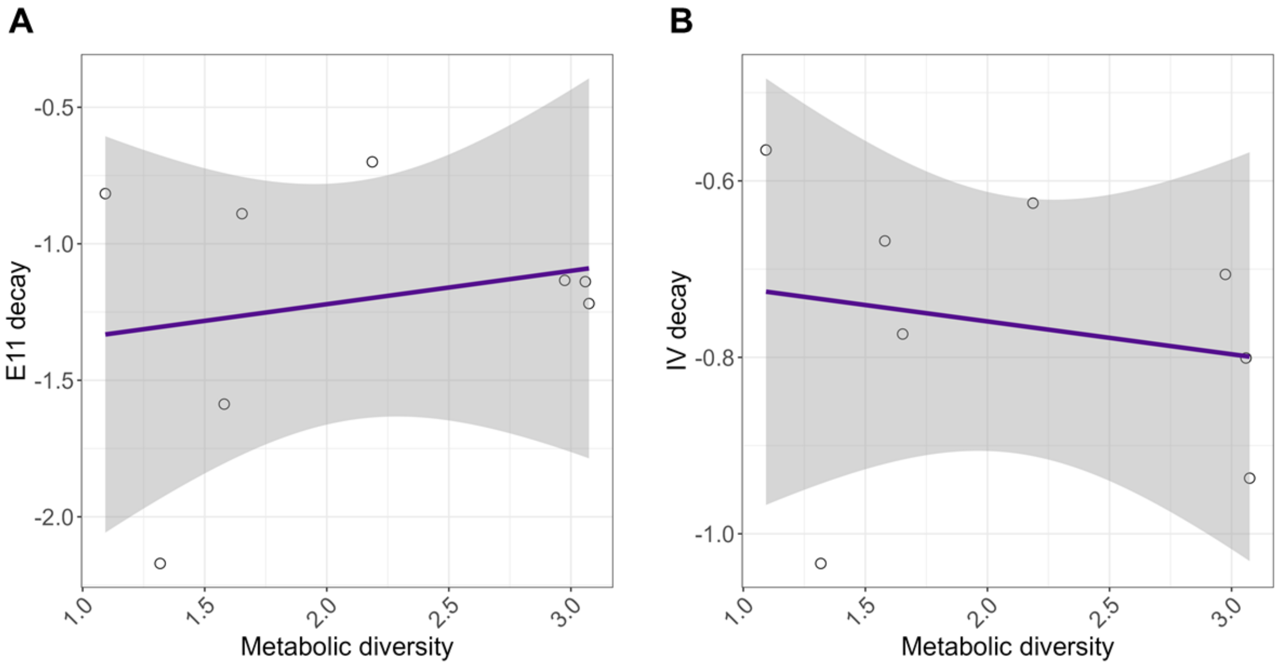


**Figure S1. Correlation between viral decay and metabolic diversity of Alpine lakes.** R studio was used to correlate viral decay values of Echovirus 11 (E11) and Influenza A virus (IAV) to Shannon diversity indexes calculated per lake using the data obtained from Biolog EcoPlates. Purple line represents the linear regression line and shaded area represents the confidence interval of the linear regression line.

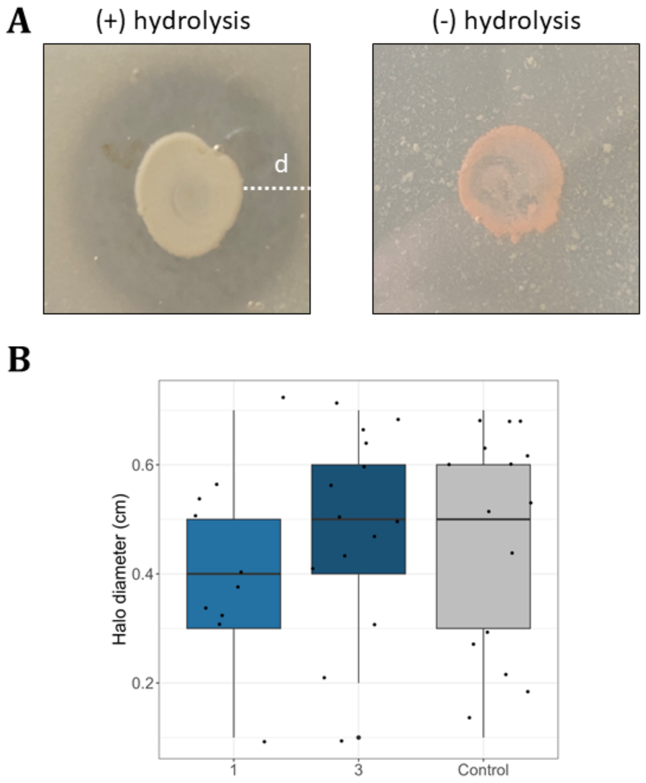


**Figure S2. Casein hydrolysis by bacteria isolates present in synthetic communities.** (A) Every isolate in the pool was grown in Skim milk agar plates as a proxy of protease production. Positive or negative hydrolysis was detected by the appearance of a halo around the colonies. (B) Casein hydrolysis of all members was compared among communities.

### Tables

#### Table S1. Date of sampling and altitude of alpine lakes sampled in this study.

| *Lake* | Date | Altitude (m) |
|---|---|---|
| *Stellisee* | 02.09.21 | 2538 |
| *Grinjisee* | 02.09.21 | 2334 |
| *Grünsee* | 02.09.21 | 2303 |
| *Mossjesee* | 02.09.21 | 2139 |
| *Leisee* | 02.09.21 | 2232 |
| *Tracouet* | 12.09.21 | 2171 |
| *Grundsee* | 17.09.21 | 1842 |
| *Guggisee* | 17.09.21 | 2007 |
| *Faflersee* | 17.09.21 | 1796 |
| *Schwarzsee* | 17.09.21 | 1861 |
| *Spilsee* | 02.10.21 | 2401 |
| *Mittelsee* | 02.10.21 | 2551 |
| *Wirbulsee* | 02.10.21 | 2658 |
| *Lengsee* | 02.10.21 | 2707 |
| *Lac des Vaux* | 08.10.21 | 2545 |
| *Small Lac des Vaux* | 08.10.21 | 2544 |
| *Oeschinensee* | 09.10.21 | 1580 |
| *Vordersee* | 15.10.21 | 2360 |
| *Petit Aletsch 1* | 15.10.21 | 2350 |
| *Petit Aletsch 2* | 15.10.21 | 2350 |
| *Petit Aletsch 3* | 15.10.21 | 2335 |
| *Marjelensee* | 15.10.21 | 2302 |
| *Lac de Salanfe* | 19.10.21 | 1925 |
| *Lac Bleu* | 25.10.21 | 2090 |

#### Table S2. Primers used in this study.

| *Target* | Primer name | Sequence (5’- 3’ ) |
|---|---|---|
| *Picoeukaryotes* | Euk345f | AAGGAAGGCAGCAGGCG |
| | Euk499r | CACCAGACTTGCCCTCYAAT |
| *Bacteria* | 27F | AGAGTTTGATCMTGGCTCAG |
| | 786R | CTACCAGGGTATCTAATC |
| *Phages* | g23_Fwd | ACWGGWCTKATYTTCGCAATG |
| | g23_Rev | AYTTYTCAACWGACCADCKACC |
| *Influenza A Matrix gene* | M30F2/08 | ATGAGYCTTYTAACCGAGGTCGAAACG |
| | M264R3/08 | TGGACAAANCGTCTACGCTGCAG |
| *Echovirus 11* | E11_VP1_F | TACCACTCGAGATCAGA |
| | E11_VP1_R | TCTCATCTGCACCATGCG |

**Table S3. Metabolic diversity indexes.**

| ***Lake*** | **Shannon** |
|---|---|
| *Faflersee* | 2.440455 |
| *Grindjisee* | 2.724327 |
| *Grundsee* | 2.816062 |
| *Grunsee* | 2.671716 |
| *Guggisee* | 2.974653 |
| *Lac Bleu* | 3.073953 |
| *Lac De Vaux* | 1.696464 |
| *Leisee* | 0.986421 |
| *Lengsee* | 2.08933 |
| *Marjelensee* | 1.579767 |
| *Mittelsee* | 2.186436 |
| *Mossjesee* | 1.317553 |
| *Petit Aletsch 1* | 3.066021 |
| *Petit Aletsch 2* | 2.562114 |
| *Petit Aletsch 3* | 2.895099 |
| *Salanfe* | 1.093009 |
| *Schwarzsee* | 3.058899 |
| *Small Lac de Vaux* | 1.652769 |
| *Spilsee* | 1.606728 |
| *Stellisee* | 0.692476 |
| *Tracouet* | 2.542304 |
| *Vordersee* | 2.210877 |
| *Wirbulsee* | 2.33329 |
